# A near gap-free haplotype-resolved genome assembly of *Zoysia japonica* uncovers intra-subgenomic gene expression and regulatory variation

**DOI:** 10.1101/2025.09.22.676383

**Authors:** Sae Hyun Lee, Preethi Purushotham, Ambika Chandra, Murukarthick Jayakodi

## Abstract

Zoysiagrass is a warm-season allotetraploid turfgrass valued for stress tolerance, but high heterozygosity and polyploidy have impeded accurate genome assembly. Here, a near gap-free, allele-aware, fully phased genome of *Zoysia japonica* cv. Palisades was assembled using PacBio HiFi and Hi-C data. The assembly resolves 20 chromosome pairs (317.2 and 317.0 Mb for haplotypes 1 and 2) with telomeric repeats at both ends of all chromosomes, attains 98.3% BUSCO completeness and 99% completeness by Merqury, and comprises approximately 50% repetitive DNA and about 40,000 protein-coding genes per haplotype. Comparative analysis identified roughly 2.9 million SNPs and numerous structural variants. Hemizygous genes were widespread (4,863 in hap-1 and 4,519 in hap-2), enriched within insertion regions, highly divergent regions, and haplotype-unique sequences, and showed elevated expression relative to homozygous genes. Among 539 one-to-one coding-identical homologs, 29 exhibited haplotype-biased expression associated with upstream promoter variants. This phased genome enables marker development, genetic mapping, and trait dissection in heterozygous polyploids.

## Main

Zoysiagrass (*Zoysia* spp. Willd.) is a perennial warm-season turfgrass widely used in lawns, sports fields, golf courses, and urban landscapes across the southern United States and East Asia, including China, Japan, and Korea (Patton et al., 2017). Species in this genus are allotetraploids (2n = 4x = 40) belonging to the Poaceae family (subfamily Chloridoideae), with genome sizes of 300–400 Mb (Tanaka et al., 2016). Their genetic variability in drought, heat, cold, shade, and salinity makes zoysiagrass an important model for studying plant responses to multiple abiotic stresses. Among these species, *Zoysia japonica* Steud. and Z. *matrella* (L.) Merr. are most extensively used in breeding programs (Chandra et al., 2017). However, its high heterozygosity resulting from outcrossing complicates genome assembly, often yielding mosaic, also referred to as haplotype-collapsed genome assemblies with mixed homologous chromosomes (Shen et al., 2025; Tanaka et al., 2016). Such genome assemblies lack the resolution to distinguish homologous and homeologous genes with distinct allelic and regulatory features, obscure haplotype-specific alleles, also known as hemizygosity, and limit the detection of phased DNA markers essential for breeding. Further, such assemblies restrict the accuracy of genetic mapping and the dissection of complex quantitative traits in these species. To overcome these limitations and accelerate breeding research, we report here a near gap-free, fully chromosome-phased genome assembly of the *Z. japonica* cultivar ‘Palisades,’ a widely used coarse-textured zoysiagrass in the southern United States.

## Results and Discussion

First, a fully phased *de novo* contig assembly was generated using PacBio HiFi reads and Hi-C data (**Table S1**) with hifiasm (Cheng et al., 2021), and subsequently scaffolded to the chromosome-scale using the allele-aware HapHiC (Zeng et al., 2024). Chromosome numbers were assigned based on alignment with the genetic map (Wang et al., 2015). The high quality of the assembly was supported by clear diagonal Hi-C contact matrices (**Figure S1**) and strong concordance with the genetic map (**Figure S2**). The final assembly comprised 20 chromosome pairs, totaling 317.2 Mb for haplotype 1 (hap-1) and 317.0 Mb for haplotype 2 (hap-2) (**Table S2, Figure 1A**). All chromosomes were gapless, except for five chromosomes in hap-1 and one in hap-2 (**Table S2**). Telomeric repeats were detected at both ends of all chromosomes (**Figure S3**), indicating telomere-to-telomere continuity in the genome. Gene space assessment by BUSCO identified 98.3% of conserved Poales orthologs genes, and assembly evaluation by Merqury estimated 99% completeness with a consensus quality of 76.5 (**Table S3**), confirming the high accuracy of the assembly. Because the progenitors of *Z. japonica* remain unidentified, we attempted subgenome assignment using *k*-mer profiling by SubPhaser (Jia et al., 2022) and phylogenetic approaches; however, both methods were inconclusive, indicating the requirement for genomic information from at least one diploid progenitor. Hap-1 and hap-2 showed overall collinearity with the previous haplotype-collapsed *Z. japonica* cultivar ‘Compadre’ assembly (**Figure S4**), but each haplotype of Palisades exhibited a distinct number structural sequence differences compared to Compadre (**Table S4**), underscoring the limitations of haplotype-collapsed assemblies in identifying true genetic variants and emphasizing the necessity of phased assemblies to accurately capture genetic diversity between zoysiagrass genotypes. Consistent with previous studies (Tanaka et al., 2016; Wang et al., 2015), both haplotypes also exhibited strong collinearity with rice (*Oryza sativa* IRGSP-1.0) and sorghum (*Sorghum bicolor* NCBIv3) (**Figure S5**), highlighting the utility of *Z. japonica* for cross-species comparative genomics and translational research.

**Figure 1.**
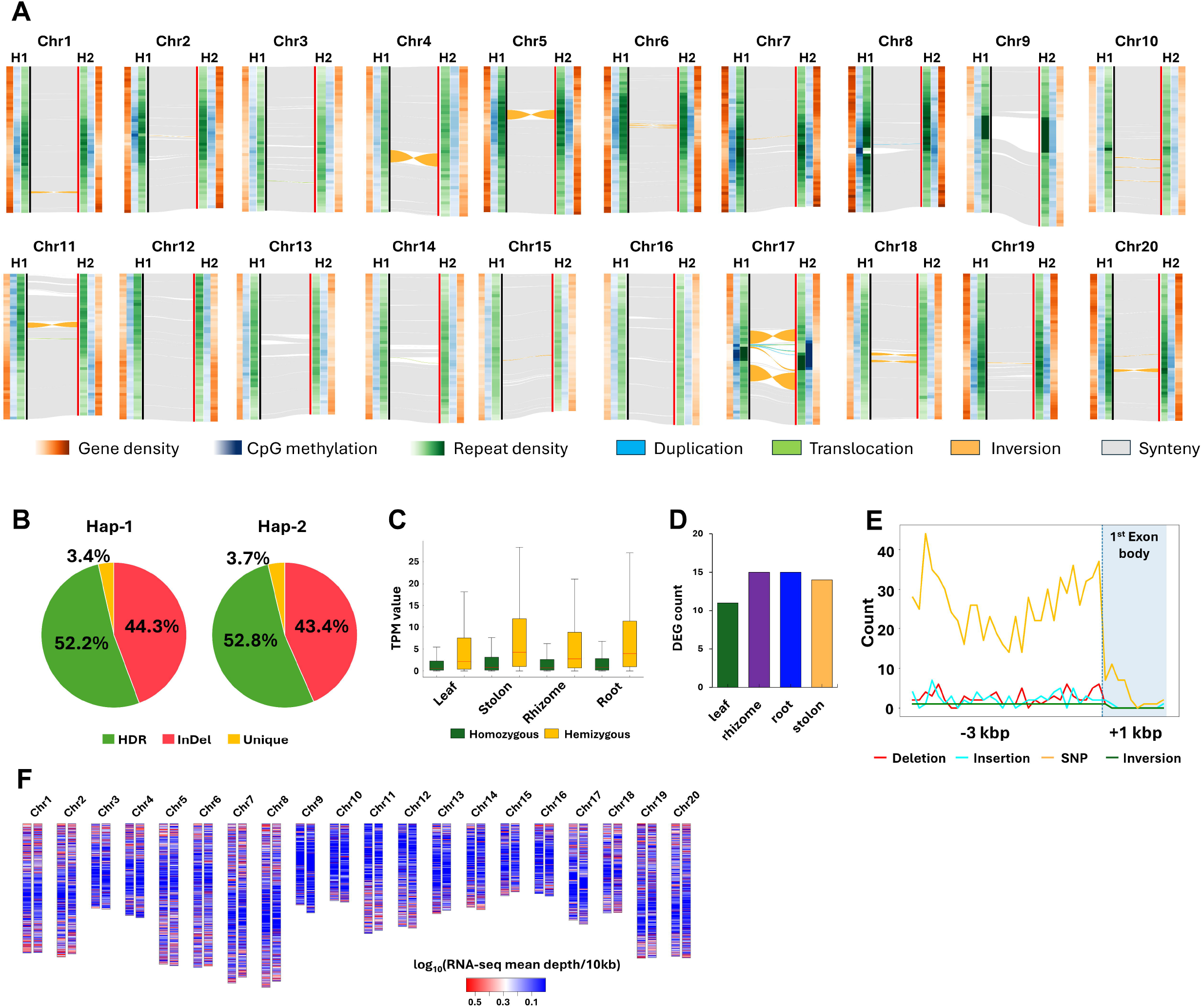
Phased homologous chromosomes and regulatory landscapes in *Z. japonica*. (A) Whole-genome collinearity between haplotypes. Heatmaps show gene density (orange), CpG methylation (navy), and repeat density (green). Syntenic regions are in gray, with duplication, translocation, and inversion events shown in blue, green, and orange, respectively. (B) Sequence features overlapping with hemizygous genes. (C) Transcript abundance (TPM) of homozygous and hemizygous genes across four tissues. Boxes indicate the interquartile range (Q1–Q3), horizontal lines mark the median, and whiskers extend to 1.5× the IQR. (D) *Cis*-regulatory differentially expressed genes (DEGs) between haplotypes across four tissues. (E) Sequence variants within 3-kb upstream regions of DEGs. (F) Expression differences between hap-1 and hap-2.

Annotation of repetitive sequences accounted for approximately 50% of each haplotype (158 Mb in hap-1 and 156 Mb in hap-2) (**Table S5**). Gene prediction, integrating short- and long-read transcriptome data (**Table S6)** and protein homology, after our rigorous manual curation (**Figure S6**), identified 40,106 and 40,223 protein-coding genes in hap-1 and hap-2, respectively (**Table S7**), with 96% BUSCO completeness. Comparison of hap-1 and hap-2 using SyRI (Goel et al., 2019) revealed extensive collinearity (∼250 Mb per haplotype) and substantial haplotype-specific sequences (142 Mb in hap-1 and 141 Mb in hap-2) (**Figure 1A**). Notably, we identified approximately 2.9 million SNPs and numerous structural variants (SVs), including 188,877 insertions, 196,125 deletions, 46 inversions, and 52 translocations (**Table S8, Figure 1A**). Most SVs were <10 kb, with only 4 inversions exceeding 1 Mb (**Table S8**). PCR validation of five randomly selected SVs confirmed their authenticity and precise breakpoints (**Figure S7, Table S9**), underlining the accuracy of the assembly and suitability of developing phased DNA markers.

A significant number of haplotype-specific hemizygous genes were identified at the intra-subgenomic level between hap-1 and hap-2. In total, 4,863 (12%) and 4,519 (11%) genes were specific to hap-1 and hap-2, respectively (**Figure 1B**). Among the hap-1-specific genes, 2,154 (44.3%) were located within hap-1–specific insertions regions, 2,539 (52.2%) within highly divergent regions (HDRs; defined as regions of extensive sequence divergence flanked by syntenic regions present in both haplotypes, as identified by SyRI), and 169 (3.4%) within haplotype-unique sequences completely absent in the alternate haplotype. A similar pattern was observed in hap-2, with 1,962 (43.4%) genes positioned within insertion regions, 2,389 (52.8%) within HDRs, and 167 (3.7%) within haplotype-unique regions. These results indicate that hemizygous genes arose from SV events between hap-1 and hap-2, as well as from divergent and unique sequences likely inherited from the progenitors of each subgenome. Importantly, these haplotype-specific hemizygous genes are not merely annotation outcomes or pseudogenes; they exhibit relatively higher expression levels compared with homozygous diploid genes (**Figure 1C**). Furthermore, to explore the *cis*-regulatory landscape between hap-1 and hap-2, we identified 539 one-to-one identical homologous genes with 100% coding sequence identity and examined their expression across four tissues (leaf, stolon, rhizome, and root). Among these, 29 genes showed significant differential expression between haplotypes (**Figure 1D**). Sequence variants, including SNPs and SVs, were identified in their upstream 3-kb promoter regions (**Figure 1E**, suggesting expression difference between haplotypes due to *cis*-regulation). Collectively, these findings reveal that SVs, HDRs, haplotype-unique sequences, and *cis*-regulatory divergence underlie extensive transcriptional differences between haplotypes (**Figure F, Figure S8**).

Our haplotype-resolved genome of *Z. japonica* provides a high-quality reference to dissect intra-subgenomic variation and its regulatory consequences. The differential transcriptional landscape between haplotypes, shaped by SVs and *cis*-regulatory divergence, accentuates the importance of phased genomes in predicting phenotypic outcomes and guiding precision breeding in heterozygous polyploids. Beyond breeding applications, the extensive haplotype-specific variation uncovered here offers opportunities for precise gene discovery and for exploiting structural rearrangements and regulatory diversity in *Zoysia* improvement.

## Acknowledgements

Funding for this publication was partially provided by the USDA hatch project 7050-0 and the multi-state hatch project 7050-0 awarded to M.J. Portions of this research were conducted with the advanced computing resources and consultation provided by Texas ACM High Performance Research Computing (HPRC).

## Conflict of interest

The authors declare no conflicts of interest.

## Author contributions

M.J. and A.C. conceived the study and designed the experiment. P.P. extracted the DNA and RNA for sequencing. S. H. L. performed genome assembly, annotation, and bioinformatics analysis. M. J., S. H. L., and P. P. drafted the manuscript. All the authors have reviewed and approved the final version of the manuscript.

## Data availability statement

Genomic data supporting the findings of this study are available in the NCBI database (https://www.ncbi.nlm.nih.gov/sra) under the BioProject accession PRJNA1314844. The genome and annotation files are available in the FigShare database (10.6084/m9.figshare.30131875).

## Notes

### Competing Interest Statement

The authors have declared no competing interest.

