## Supplementary materials for "A near gap-free haplotype-resolved genome assembly of *Zoysia japonica* uncovers intra-subgenomic gene expression and regulatory variation"

**DNA extraction and sequencing**

Young, fresh leaf tissues were ground in liquid nitrogen, and high-molecular-weight (HMW) DNA was extracted using Carlson buffer (Carlson *et al.*, 1991). DNA was further purified using the Genomic-tip 100/G kit (Qiagen, Hilden, Germany) following the manufacturer’s protocol. Purified DNA was sequenced using the PacBio Revio System (Pacific Biosciences, Menlo Park, CA, USA) to generate HiFi circular consensus reads. HiFi BAM files were exported to the FASTQ format using SMRT Link v25.2 (bam2fastq) (https://www.pacb.com/support/software-downloads/) (**Table S1**). An Omni-C library was prepared using the Omni-C kit (Dovetail Genomics, Scotts Valley, CA, USA) and sequenced on the NovaSeq 6000 platform (Illumina, San Diego, CA, USA) according to the manufacturer’s instructions (**Table S1**).

**Genome assembly, evaluation and repeat annotation**

HiFi reads were assembled using hifiasm v0.24.0-r702 (Cheng *et al.*, 2021) in Hi-C mode using paired Omni-C reads. Omni-C reads were aligned to the resulting phased contigs using BWA-MEM v0.7.17 (Li, 2013), and primary scaffolding was performed using the HapHiC pipeline v1.0.6 (Zeng *et al.*, 2024). Scaffolds were manually curated using the Juicebox Assembly Tools v3.1.4 (Durand *et al.*, 2016). Scaffolding accuracy was further validated by anchoring to a previously published genetic map (Wang *et al.*, 2015). Assembly completeness and consensus quality were assessed using Merqury v1.3 (*k*-mer–based completeness and QV) (Rhie *et al.*, 2020). Gene space completeness was evaluated with BUSCO v5.7.1 and compleasm program against the poales_odb10 dataset (Huang and Li, 2023; Manni *et al.*, 2021; Simão *et al.*, 2015). Telomeric repeats were identified by scanning for the canonical motif (TTTAGGG). Repetitive elements were annotated with EDTA v2.1.0. (Ou *et al.*, 2019).

**Transcriptome sequencing**

Total RNA was extracted from leaf, stolon, rhizome, and root tissues using the Macherey-Nagel NucleoSpin RNA Plant Mini Kit according to the manufacturer’s protocol. All extractions included the DNase step and were quality checked using a Bioanalyzer 2100 (Agilent). RNA was sequenced on the NovaSeq 6000 and adapter-trimmed with Trim Galore v0.6.10 (Cutadapt v4.6; Q≥20, minimum length ≥20 bp) (**Table S6**) (Martin, 2011). For long-read MAS-Seq, cDNA libraries were prepared with the Kinnex full-length cDNA kit and sequenced on the Revio System (Pacific Biosciences) (**Table S6**). MAS-Seq data were processed using SMRT Link v25.2.0 (isoseq refine; isoseq cluster2). Barcode demultiplexing and adapter removal were performed using Lima v2.6.0 (https://github.com/PacificBiosciences/pbbioconda). High-quality, full-length non-chimeric (FLNC) reads were identified using isoseq refine, and consensus isoforms were obtained using isoseq cluster2 (Al’Khafaji *et al.*, 2024) (https://github.com/PacificBiosciences/pbbioconda).

**Gene prediction**

For gene prediction, the RNA-seq data generated in this study and previous studies (**Table S10**) were included. Short-read RNA-seq data were aligned to the assembled genome using HISAT2 v2.2.1 (Kim *et al.*, 2019). To provide protein homology support, protein sequences from closely related species, including *Eremochloa ophiuroide* (Wang *et al.*, 2021), *Oryza sativa* (Kawahara *et al.*, 2013), *Sorghum bicolor* (McCormick *et al.*, 2018) , *Triticum aestivum* (Consortium (IWGSC) *et al.*, 2018), and *Zea mays* (B73, NAM-5.0) (Hufford *et al.*, 2021) were used. Gene prediction was performed using BRAKER3 v3.0.8 (Gabriel *et al.*, 2024), which integrates RNA-seq and protein evidence. Full-length MAS-Seq reads were mapped to the genome using minimap2 v2.26 (Li, 2018). Redundant isoforms were collapsed to retain the longest representative transcript models using cDNA_Cupcake v29.0 (<https://github.com/Magdoll/cDNA_Cupcake>). Coding regions were predicted from non-redundant isoforms using GeneMarkS-T v5.1 (Tang *et al.*, 2015). BRAKER3 predictions and GeneMarkS-T models were subsequently integrated with TSEBRA v1.0.3 (Gabriel *et al.*, 2021; Tang *et al.*, 2015) to produce consensus annotations. MAS-Seq–based predictions were compared with the consensus results to identify and correct potential misannotations (**Figure S6**). The final gene set was refined by filtering out transposable element (TE)-related genes based on overlap with repeat annotations and protein domain searches using InterProScan v5.59-91.0 (Jones *et al.*, 2014) and eggNOG-mapper v2.1.12 (Cantalapiedra *et al.*, 2021). Long non-coding RNAs were predicted using FEELnc v0.2 (Wucher *et al.*, 2017) and were excluded from the protein-coding gene set. Genes with coding sequences shorter than 300 bp and lacking functional domains were discarded.

**Variant calling**

Phased haploid genomes were aligned using mm2plus v1.0 (Chandra *et al.*, 2024). Sequence variations, including single-nucleotide polymorphisms (SNPs) and structural variants (SVs), were identified using SyRI v1.7.0 (Goel *et al.*, 2019) with default parameters. SVs were visualized using plotsr v1.1.0 (Goel *et al.*, 2019; Goel and Schneeberger, 2022).

**Identification of homologous genes between haploid genomes**

Transcript sequences from haplotype 1 and haplotype 2 were compared to identify homologous gene pairs. BLASTN (Camacho *et al.*, 2009) searches were conducted (e-value ≤ 1e–5). For each transcript, the top-scoring hit was retained, and reciprocal best hits with complete sequence identity and matching chromosomal assignment were defined as homologous genes for downstream analyses.

**Expression quantification and CpG methylation profiling**

RNA-seq reads were aligned to the haplotype-resolved genome assemblies using HISAT2 v2.2.1 (Kim *et al.*, 2019). Following alignment, transcript abundance was quantified in terms of transcripts per million (TPM) using TPMCalculator v0.0.4 (Vera Alvarez *et al.*, 2019). CpG methylation was detected from PacBio HiFi reads using pb-CpG-tools v2.0 (https://github.com/PacificBiosciences/pb-CpG-tools).

**PCR amplification and gel electrophoresis**

PCR amplification was carried out using DreamTaq DNA polymerase (Thermo Fisher Scientific) on a Bio-Rad T100 Thermal Cycler. The cycling conditions were as follows: initial denaturation at 95 °C for 3 min, followed by 35 cycles of 95 °C for 20 s, 58 °C for 20 s, and 72 °C for 20 s, with a final extension at 72 °C for 5 min. Amplified products were resolved on a 2% agarose gel by electrophoresis at 100 V for 40 min.

**Figure legend**

**Figure S1.** Interchromosomal Hi-C contact matrix. (A) Haplotype 1. (B) Haplotype 2.

**Figure S2.** Concordance between the genetic and physical maps. The x-axis represents physical positions along the genome, and the y-axis indicates genetic distances in centimorgans (cM). (A) Haplotype 1. (B) Haplotype 2.

**Figure S3.** Genome-wide distribution of the telomeric repeat TTTAGGG. The x-axis represents physical positions, and the y-axis indicates the frequency of repeat sequences. (A) Haplotype 1. (B) Haplotype 2.

**Figure S4.** Collinearity analysis between previously reported *Compadre* (Shen *et al.,* 2025) genome and the haplotype-resolved *Palisade* genome generated in this study.

**Figure S5.** Collinearity analysis between *Sorghum bicolor* and *Oryza sativa*. (A) *Sorghum bicolor*. (B) *Oryza sativa*.

**Figure S6.** Representative cases of gene annotation model curation. Blue boxes indicate curated models, and yellow boxes represent models prior to curation. MAS-seq alignments are shown below the models. (A) An incorrectly split gene model. (B) An incorrectly merged gene model. (C) A gene model without supporting RNA evidence.

**Figure S7. Validation of SV markers.** PCR products were resolved on a 2% agarose gel. Samples are arranged from left to right as haplotype 1, ladder, and haplotype 2.

**Figure S8.** **Genome-wide heatmap of RNA expression profiles in haplotype 1 and haplotype 2.** From top to bottom, rows correspond to rhizome, stolon, and root
