## Supplementary figures and images for "A near gap-free haplotype-resolved genome assembly of *Zoysia japonica* uncovers intra-subgenomic gene expression and regulatory variation"

### Figure S1

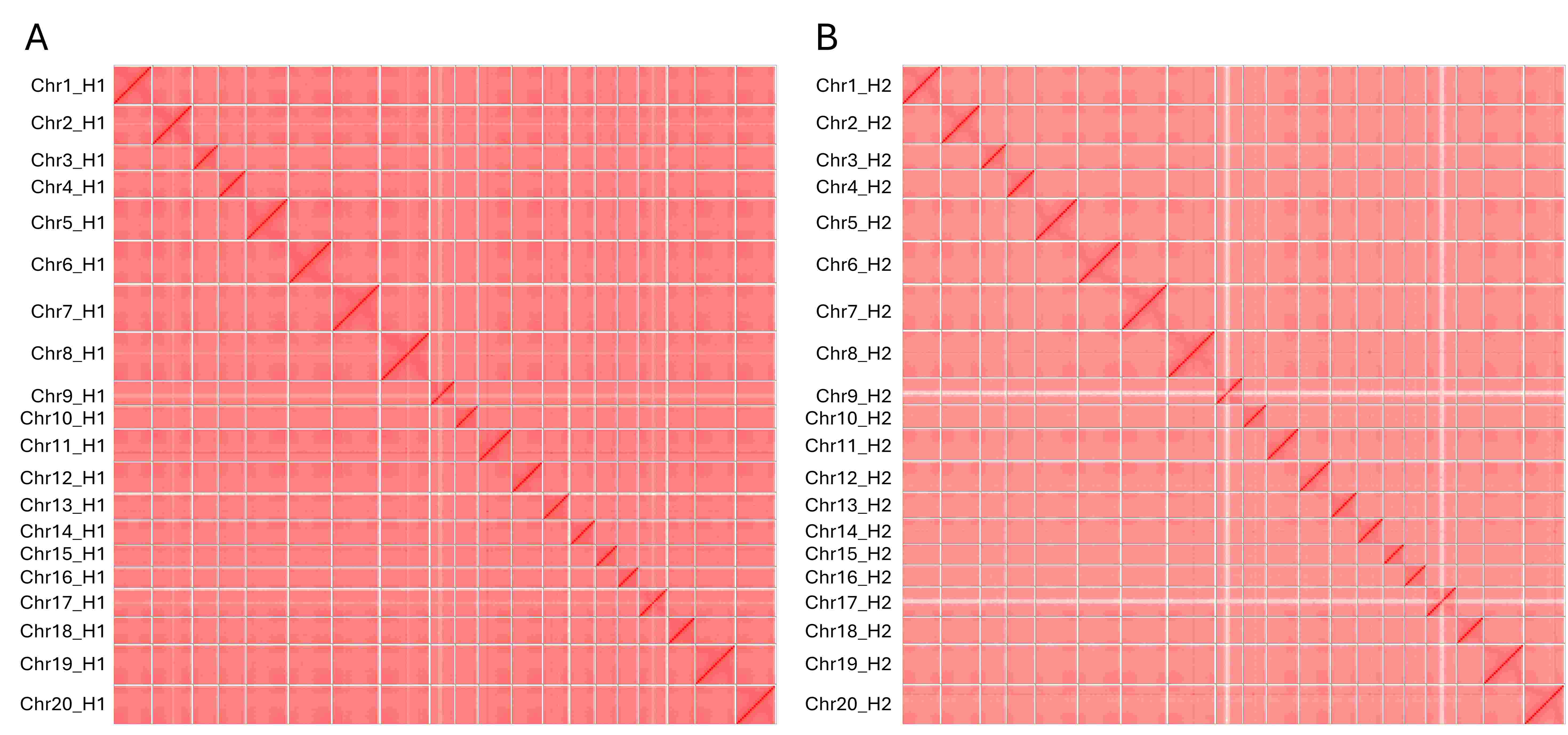

### Figure S2

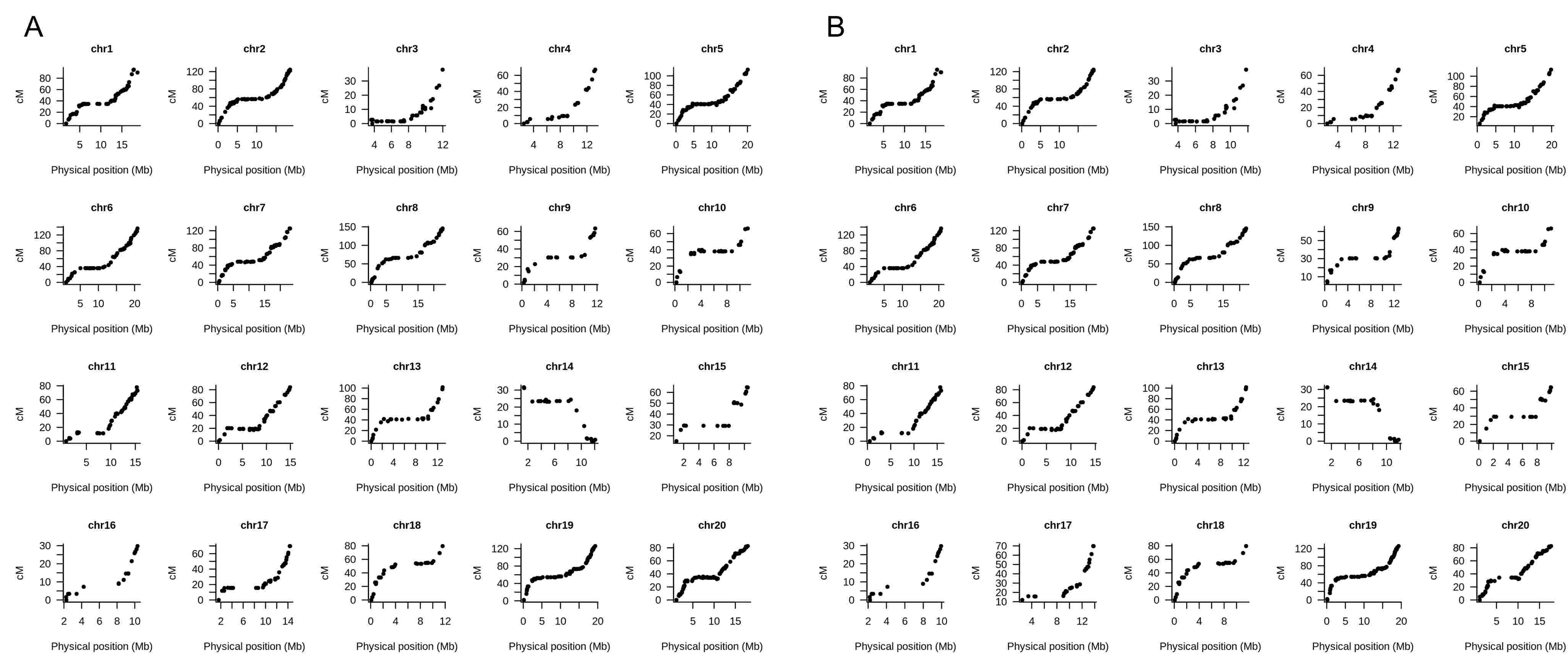

### Figure S3

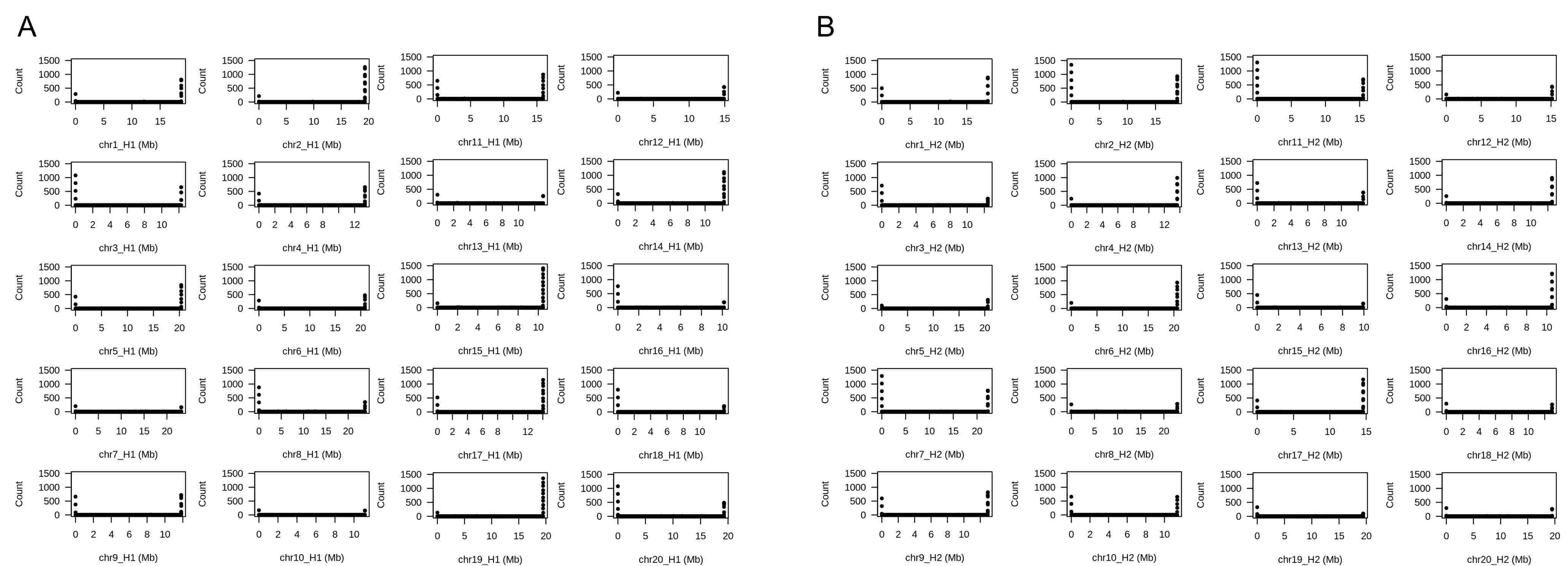

### Figure S4

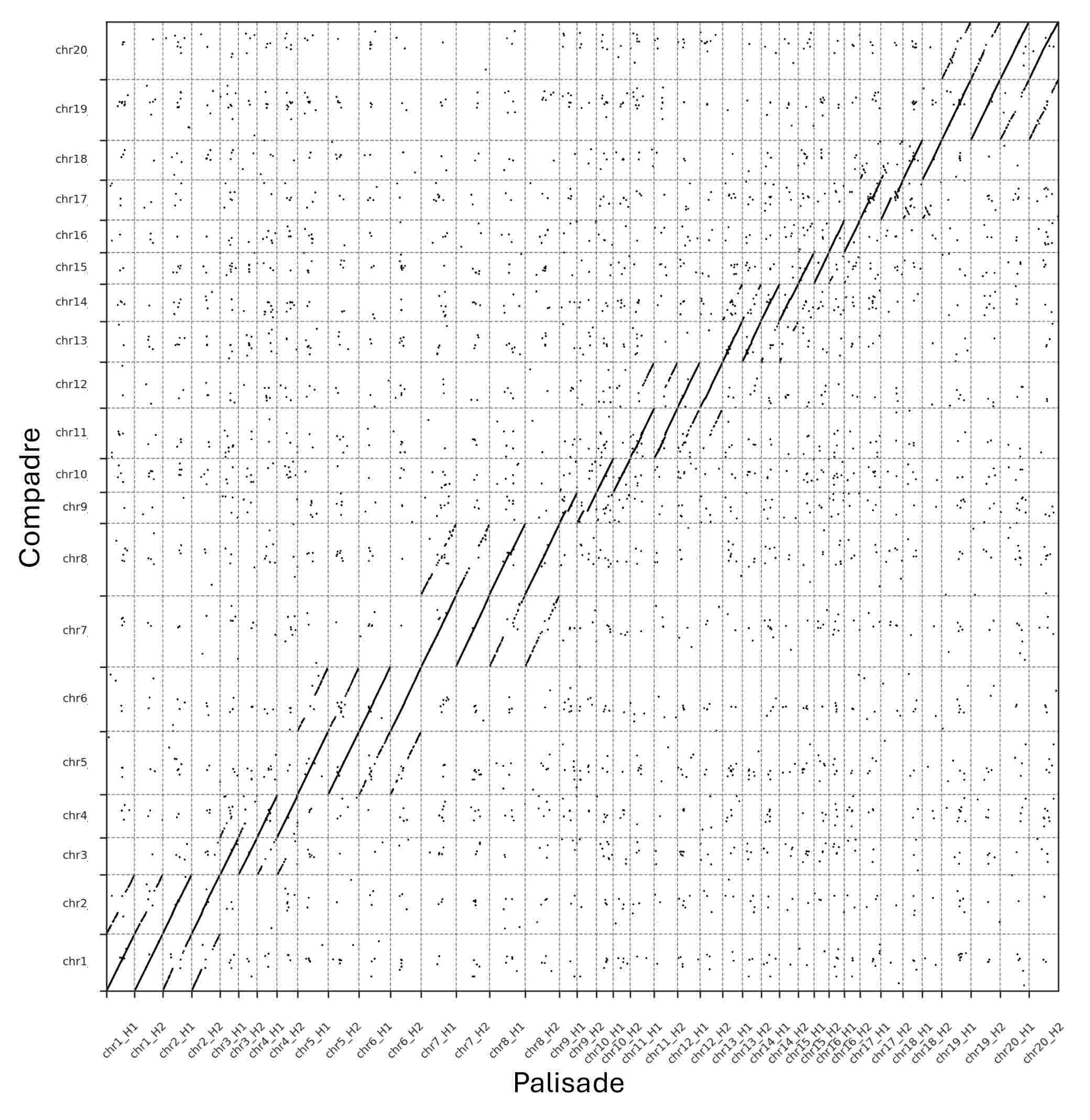

### Figure S5

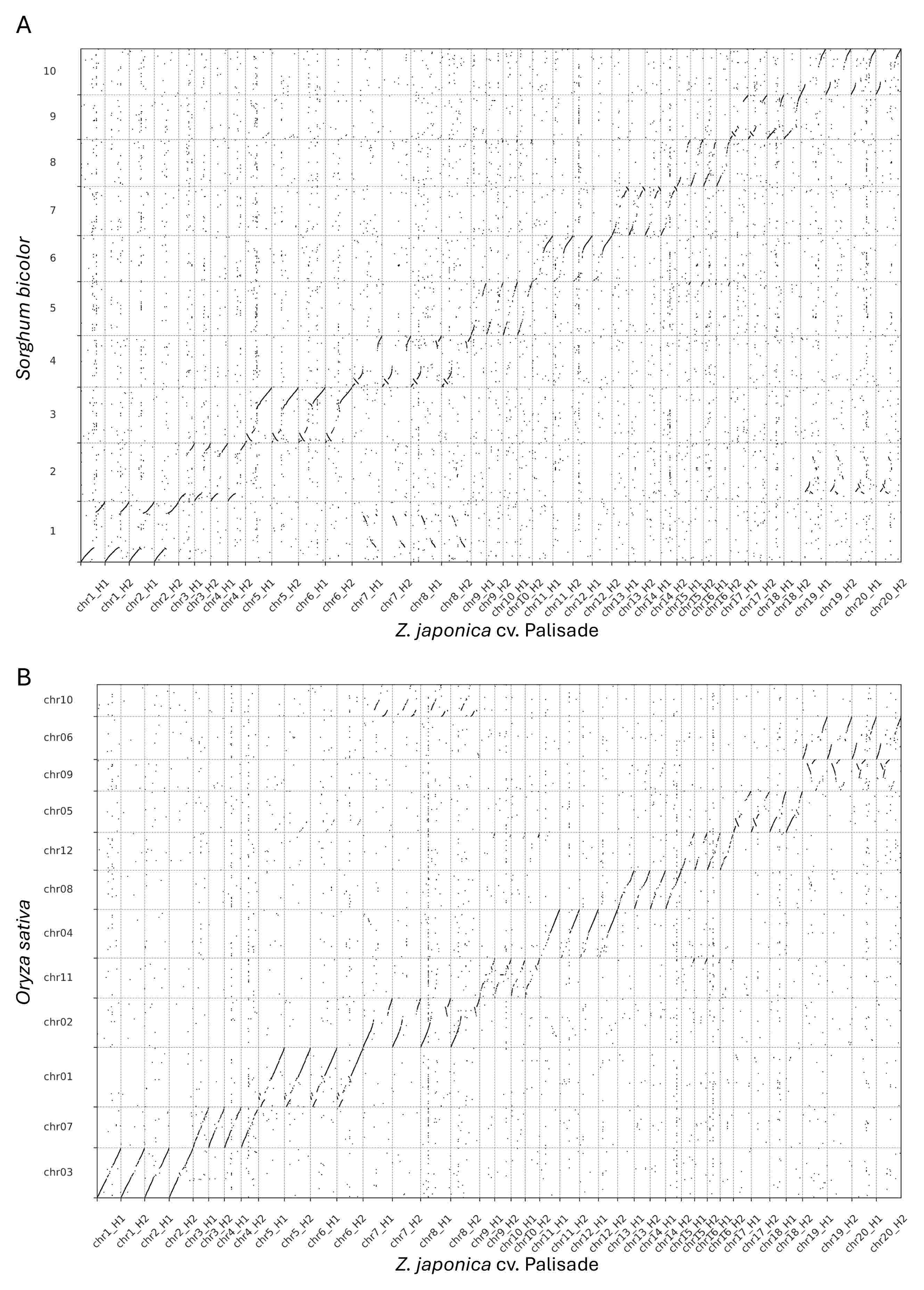

### Figure S6

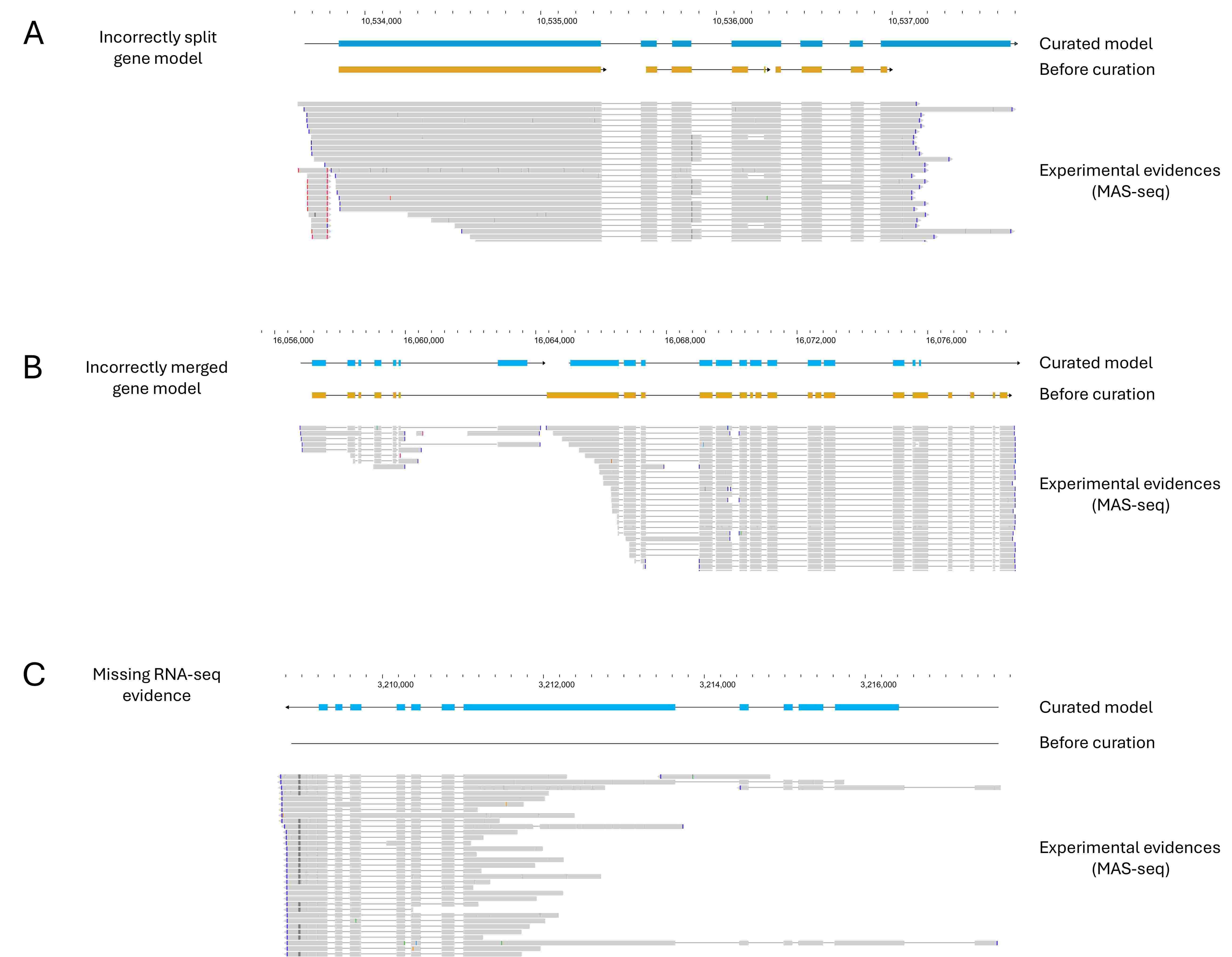

### Figure S7

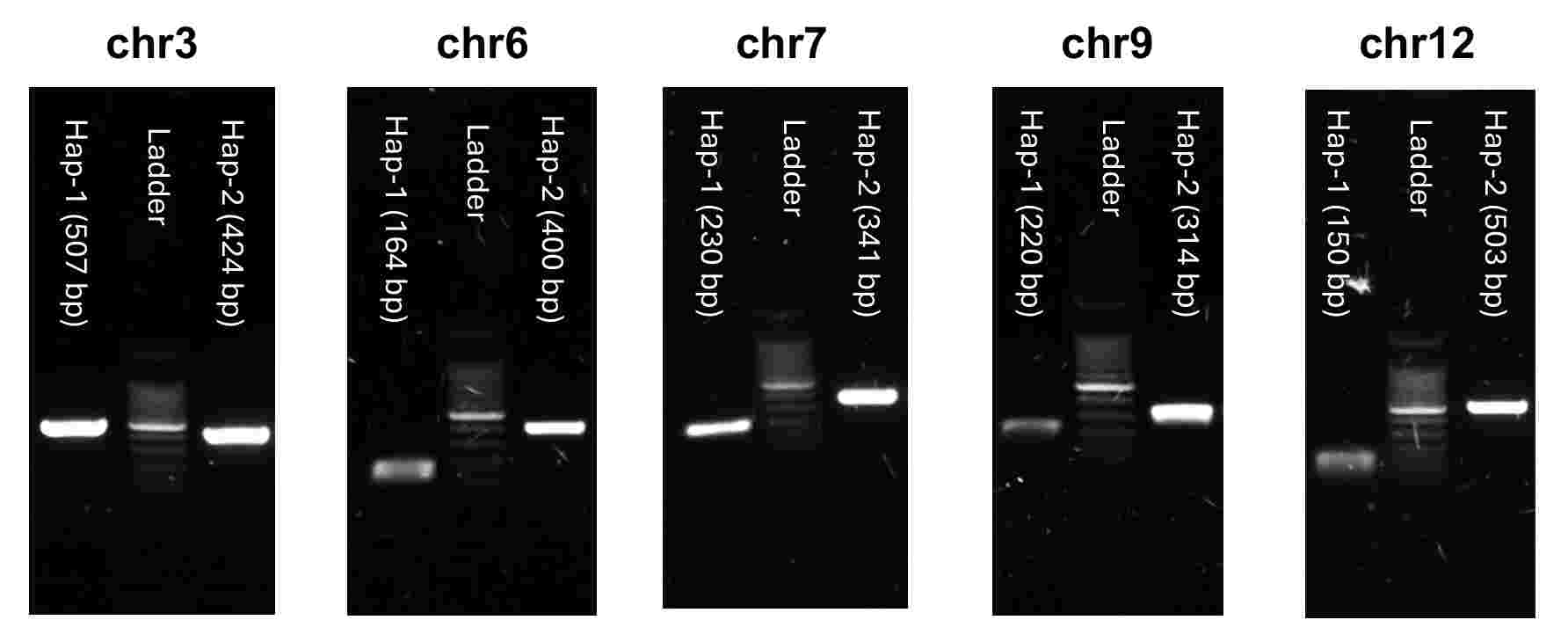

### Figure S8

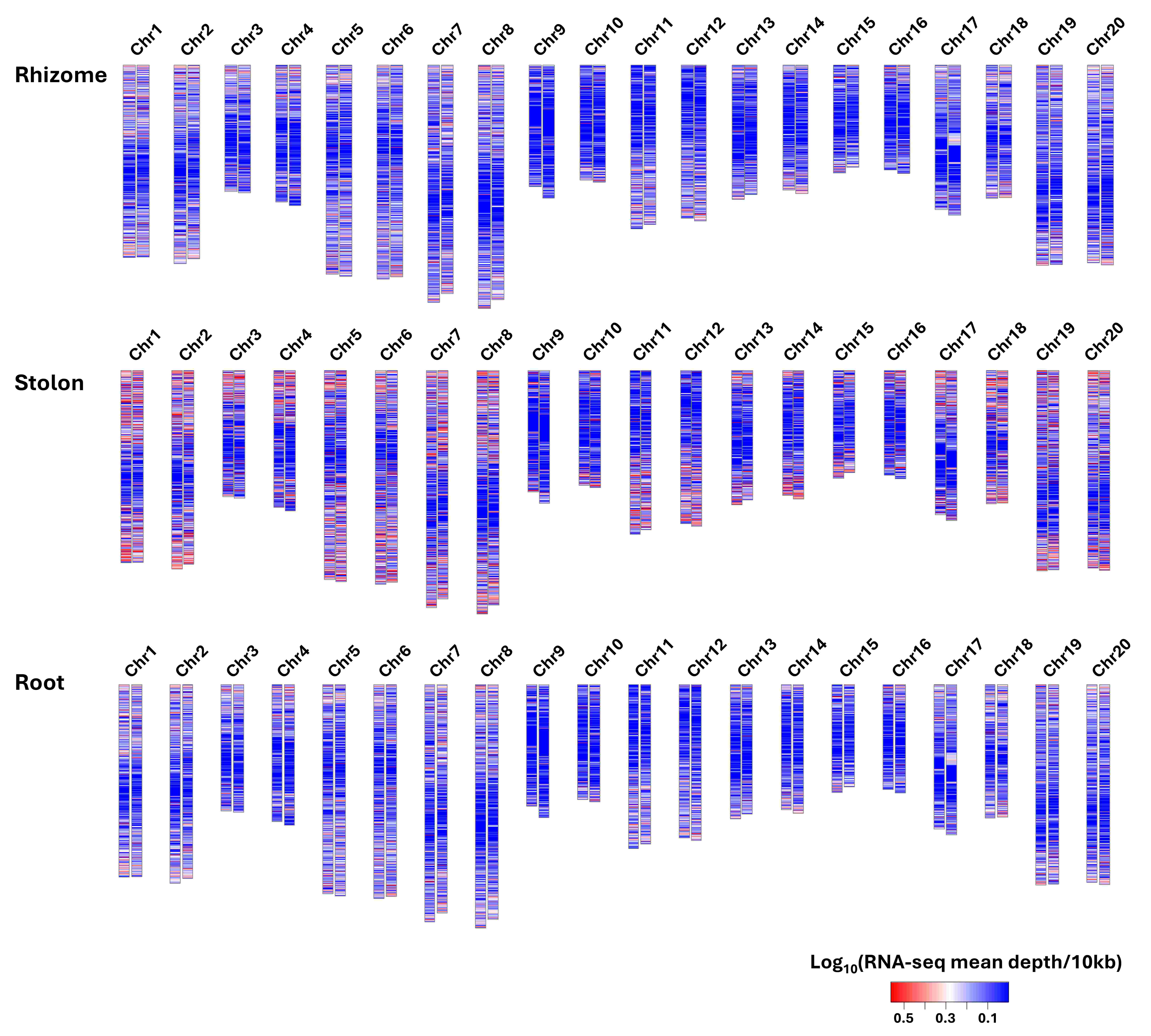
